# Reorganisation of complex ciliary flows around regenerating *Stentor coeruleus*

**DOI:** 10.1101/681908

**Authors:** Kirsty Y. Wan, Sylvia K. Hürlimann, Aidan M. Fenix, Rebecca M. McGillivary, Tatyana Makushok, Evan Burns, Janet Y. Sheung, Wallace F. Marshall

## Abstract

The phenomenon of ciliary coordination has garnered increasing attention in recent decades, with multiple theories accounting for its emergence in different contexts. The heterotrich ciliate *Stentor coeruleus* is a unicellular organism which boasts a number of features which present unrivalled opportunities for biophysical studies of cilia coordination. With their cerulean colour and distinctive morphology, these large protists possess a characteristic differentiation between cortical rows of short body cilia used for swimming, and an anterior ring structure of fused oral cilia forming a membranellar band. The oral cilia beat metachronously to produce strong feeding currents. In addition to this complex body plan, *Stentor* have remarkable regenerative capabilities. Minute fragments of single cells can over the period of hours or days, regenerate independently into new, proportional individuals. Certain environmental perturbations elicit a unique programmed response known as oral regeneration wherein only the membranellar band is shed and a new, ciliated oral primordium formed on the side of the body. Here, we target oral regeneration induced by sucrose-shock to reveal the complex interplay between ciliary restructuring and hydrodynamics in *Stentor*, which accompanies the complete developmental sequence from band formation, elongation, curling, and migration toward the cell anterior.

*“When the anterior part is open, one may perceive about its Edges a very lively Motion; and when the Polyps presents itself in a certain manner, it discovers, on either side of these edges of its anterior part, somewhat very much resembling the wheels of a little Mill, that move with great velocity.”*

*A. Trembley F.R.S describing the membranellar band of Stentor, Phil. Soc. Trans. Royal Society (London), 1744.*

## 1. Introduction

### 1.1. *Stentor* - the trumpet animalcule

The first recorded account of these fascinating funnel-shaped creatures was made by Abraham Trembley F.R.S, who in a letter to the president of the Royal Society [1], wrote “I have herewith the honour of transmitting to you the particulars of several observations I have made, during the course of the last summer, upon some species of very minute Water-Animals…” [2]. (In fact, at the time Trembley had misidentified these organisms as *Hydra*.) These unicellular ciliates, which are easily visible to the naked eye, are named *Stentor* for their trumpet-like morphology, in honour of a loud-voiced Greek hero who fought in the Trojan War. *Stentor* are widely-occurring in both freshwater and marine habitats, exhibiting both free-swimming and sedentary characteristics depending on environmental circumstance. Individual cells undergo dynamic and extensive shape changes due to a highly-contractile cortical structure, assuming a pear or tear-drop shape when free-swimming, contracting into a ball when disturbed, or else extending up 1mm in a rest state in which cells become attached to substrates via a posterior holdfast (Figure 1a). In the latter state, a large number of ciliated feeding organelles are visible at the anterior end of the organism, which stir the fluid collectively to produce strong feeding currents, drawing food and other particulates towards the mouth of the organism. Not only is the oral apparatus the site of a remarkable degree of ciliary coordination and highly-coupled fluid-structure interactions, it is the most differentiated cortical landmark of a *Stentor*, which can also be regenerated in response to injury or damage. The present study solicits this dramatic regenerative capability to investigate ciliary organisation and coordination - which remains an open and confounding problem in ciliary biology. Here, state-of-the-art imaging and flow field tracking is combined for the first time, to explore the extreme restructuring of cilia and extracellular flows during *Stentor* oral regeneration.

**Figure 1.**
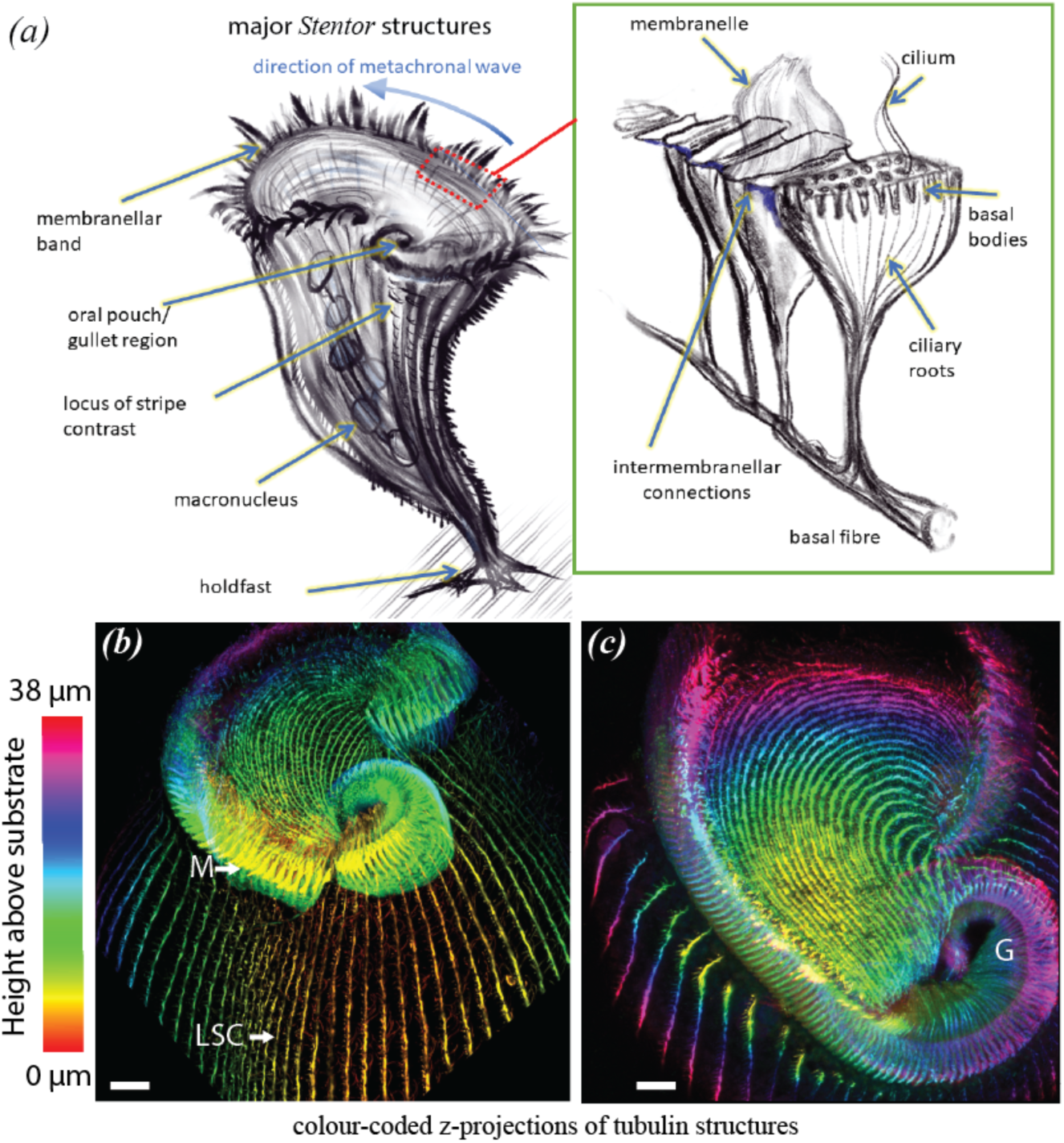
a) Schematic of a single *Stentor* cell with key features highlighted. Inset - ultrastructure of membranellar band [6] showing rows of oral cilia arranged in parallel stacks (each approximately 7.5µm x1.5µm); b) confocal immunofluorescence images showing the structure and organisation of the membranellar band (M), cortical striation patterns (notice the sharp change in width at the locus of stripe contrast: LSC), and associated rows of short body cilia, c) top view of the frontal field and gullet region (G). [Scalebars = 10 µm.]

While diverse species of *Stentor* exist, we focus exclusively on *S. coeruleus Ehrenberg* 1830 -- a widely distributed species, notable for their striking cerulean pigmentation. *S. coeruleus*, also called the *Blue Stentor* [2, 3], was favoured historically for cytological studies due to its large size and moniliform macronucleus [4], and has been the subject of recent genomic studies [5]. The conical surface of the organism bears longitudinal stripes of distinctive colouration extending from the anterior all the way down to the tapered holdfast: cortical rows of constant width, alternate with pigmented stripes of graded width – thereby breaking the patterning symmetry (Figure 1b). The pigmented bands increase in width from left to right around the cell, culminating in a special region called the locus of stripe contrast (LSC) or “ramifying zone”. This region demarcates the location where the new oral primordium and a new field of basal bodies will form.

The body cilia are inserted in rows along the clear stripes all over the body; the anterior body cilia are somewhat shorter than those at the posterior [4]. Contractile fibres called myonemes subtend each row of body cilia, and are involved in whole-body contractions. The broad anterior end of the organism is encircled by an adoral zone of external cilia that are fused into rows of transverse plates termed membranelles (*sensu* Sterki 1878). In SEM studies, Randall and Jackson showed [6] that these adoral structures were not individual large cilia but rather two or three rows of tightly-packed cilia (20-25 cilia per row), so that each membranelle maintains functional unity as an intercalated ciliary sheath (Figure 1a, inset). The entire membranellar band, also known as the *peristome*, contains around 250 such stacked membranelles; one end terminates freely, while the other end spirals into a funnel-shaped invagination or gullet (Figure 1c). The region enclosed by the membranellar band is the frontal field, which is distinguished by a change in stripe orientation – the frontal field stripes are also narrower since they originated as ventral stripes that have been shifted forward over the course of oral primordium formation (see section 2), together with the new stripes which will go on to develop in this zone.

### 1.2. Regenerative prowess of *Stentor*

Regeneration of ciliated structures provides a unique way of probing the emergence and reorganisation of ciliary interactions over developmental timescales. A hallmark of life is its ability to repair, heal wounds, or replenish lost or damaged cellular structures [7, 8]. Single cells are of special interest, for regeneration must necessarily take place entirely within the same cell. One classic example is flagellar regeneration in the biflagellate alga *Chlamydomonas*, where amputated flagella can be regrown to full length in about 1.5 hours [9, 10], whereupon biflagellar coordination also recovers [11, 12]. Early researchers demonstrated the versatility of *Stentor* as a model system for studies of regeneration, made possible by their large size and manoeuvrability, as well as remarkable regenerative capabilities [13, 14]. Questioning the limits of cellular indivisibility led the likes of F.R. Lillie and T.H. Morgan [15, 16] to conduct dissection experiments on *Stentor* to determine the minimal ingredients for regeneration. They found that smallest of fragments capable of regenerating into new individuals that would go on to divide and multiply, were spheroids of only <100 µm in diameter. Nucleated fragments irrespective of whether they originate from the head or tail, are all capable of full regeneration.

Here, we study one particular developmental sequence of events which features consistently during *Stentor* regeneration, in which a new membranellar band is formed *de novo* (Figure 2a-g). This characteristic program is followed routinely by cells undertaking physiological cell division to develop a second oral primordium [17], and also by cells that have been induced by diverse treatments (such as dissections, grafting and chemical exposure [18]) to shed and subsequently replace an existing primordium and associated oral structures. Initiation of regeneration requires the presence of at least one macronuclear node, some cytoplasmic material, as well as a piece of the microtubule cortex. Selective inhibition studies have shown that DNA-dependent RNA synthesis is required to supply the new protein required during intermediate stages of regeneration, but not during the later stages. Depending on temperature, medium, and other unknown factors [19], the total regeneration time ranges between 8-10 hours at 20-25 °C. Wherever successful, regeneration proceeds through the following major stages (Figure 2a): i) within the first hour a rift forms at the site of the new primordium; ii) the first cilia sprout, reaching ∼ 0.5 µm, new basal bodies form in the vicinity of existing somatic cilia, membranellar cilia are distributed randomly, and beating is uncoordinated at this stage; iii) at 3-4 hours proliferation of basal bodies continues and the new primordium attains the definitive width of 12 µm, cilia approach full length though shorter cilia are still evident, and some evidence of coordinated ciliary activity is visible in SEM sections [20]; iv) cilia assemble into membranelles and the membranellar band structure continues to lengthen, the moniliform nucleus condenses; v) at 6.5-7 hours, significant membranellar band migration towards the anterior of the organism occurs, vi) finally, after ∼8 hours, the newly formed membranellar band curls all the way around the anterior end, and the nucleus renodulates.

**Figure 2:**
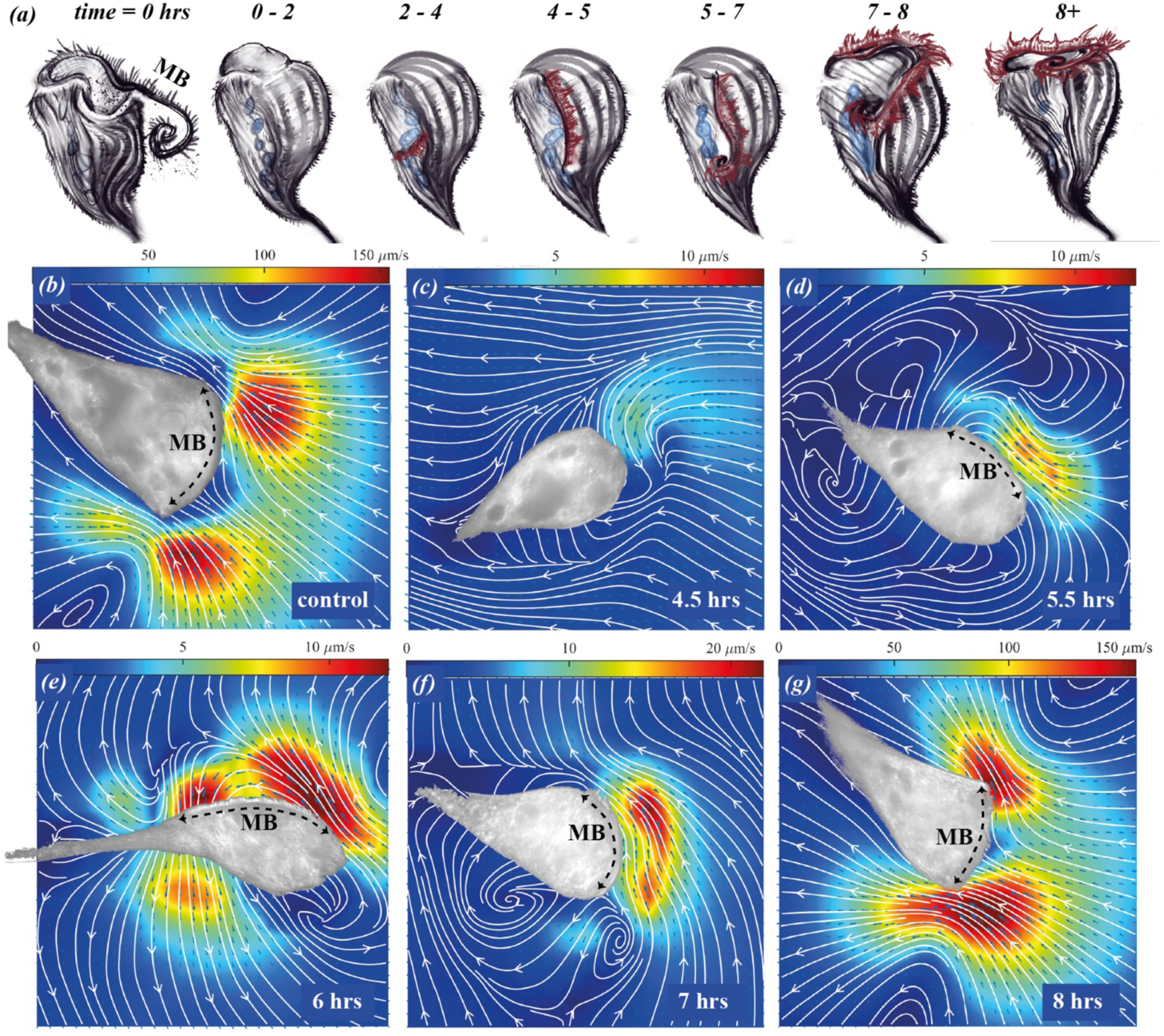
a) Chronology of membranellar restructuring during induced oral regeneration in *Stentor coeruleus*: at time 0 the existing membranellar band was induced to shed by sucrose shock, after 1-2 hrs a rift opens in the vicinity of the indifferent zone, between 2-5 hrs membranellar cilia sprout, lengthen, and rearrange themselves into rows of stacked membranelles, after 5 hrs, the membranellar band undergoes elongation and gradual migration to finally assume a nearly-circular structure at the anterior end; b)-g) PIV measurements of the extracellular flow fields associated with the regenerating membranellar band for a control cell, and at the indicated times post-shock for different regenerating cells. [Colormaps indicate flow speed. Masked, brightfield images of the adhered organisms have been overlaid on top of the flow maps. MB demarcates the location of the membranellar band - wherever it is clearly identifiable.]

For this study, we regenerated ∼80 *Stentor* cells, and can confirm all qualitative morphological features of the above staging. Oral regeneration in *S. coeruleus* was induced by sucrose shock, with slight modifications from published protocols [21, 5], which were based on historical works (see supp. materials). Briefly, *Stentor* were exposed to a 10% sucrose solution for 2-3 minutes, which elicited synchronous shedding of the membranellar band in >80% of the cases (supp. Materials). Cells were then left to recover in fresh medium, and extracellular flows and ciliary activity accompanying regeneration were analysed on a cell-by-cell basis.

## 2. Reorganisation of ciliary flows during regeneration

In order to measure the flow fields around *Stentor* at high spatiotemporal resolution, we developed a simple but effective protocol for preventing cell body motion, adapted from conventional cell attachment techniques. Glass-bottom petri dishes used for imaging were pre-treated with Poly-D-Lysine to encourage surface adherence of *Stentor* (supp. Materials). Contrary to many other methods trialled previously, this particular surface attachment protocol had minimal effect on cell viability, allowing for live-cell imaging for extended periods of time, and did not adversely alter the oral regeneration dynamics. The boundary conditions consisted of a free surface (a large open droplet), and a flat solid substrate. To obtain flow fields around the regenerating organisms, we seeded the medium with passive tracers (1 µm polystyrene beads) and performed Particle Image Velocimetry (PIV) [22]. The body cilia remained motile but beat only intermittently throughout the oral regeneration process, and were capable of producing large-scale flows. This unsteady nature of ciliary activity is not fully-understood, but also pertains to non-adhered organisms (supp. Figure S1). Here, in order to isolate flow contributions due to membranellar cilia only, we restrict our discussion and analysis to recordings which do not show excessive body cilia motion or body contractions (supp. Figure S2), and in which flows remained steady over the entirety of each 30s recording.

### 2.1. Early regeneration and a linear membranellar band (0-5 hrs post-sucrose shock)

Under brightfield microscopy, the preliminary stages of regeneration were particularly difficult to discern in the substrate-adhered organisms -- it is not possible to see the newly-formed rift and elongating membranellar band until after a substantial structure has been formed. Sucrose-induced shedding rarely removes 100% of the membranelles -- in most cases a small number of these will remain deep inside the old gullet, destined to merge with the newly formed membranellar band once the latter migrates successfully all the way to the anterior. In these cases, the old membranelles can produce weak flows towards the gullet (Figure 2c). The new membranellar cilia reach a perceptible length at 4-5 hours, forming a linear band on the side of the organism.

### 2.2. Membranellar band growth and reorientation (5-7 hrs post-sucrose shock)

As the membranellar cilia continue to grow, ciliary coordination increases concomitantly (see section 3). The new membranellar band is usually conspicuous under DIC microscopy at 5 hours after sucrose shock, and associated with localised flows that are directed posteriorly and largely parallel to the cell surface (Figure 2d, supp. Video 1). Over the next hour, the membranellar cilia proceed to lengthen and reorganise [20], producing flows which exhibit a greater perpendicular component that is directed away from the cell body (Figure 2e). Flow magnitudes remain low during this period, on the order of 10 µm/s. After ∼7 hours, the band structure migrates upwards and the new gullet begins to form, the cell anterior becomes more conical and distortion of streamlines is observed (Figure 2f).

### 2.3. Emergence of the feeding vortex and completion of regeneration (>7 hrs post-sucrose shock)

As the arc of the developing membranellar band continues to move upward, the two ends eventually approach each other and the enclosed stripes bend into arcs in the frontal field (Figures 1, 2a). Finally, a characteristic double-vortex pattern emerges – as in the control organisms (Figures 2b, g). In fully-regenerated *Stentor*, the membranelles exhibit robust phase coordination and metachronal waves (MCW), which results in a sharp transition to faster flows (order 100 µm/s) and appearance of the “alimentary vortex” *sensu* Maupas [2]. Structurally similar flow fields were also measured in *Stentor* that have not been adhered to a substrate, but rather were attached spontaneously by their posterior holdfast (supp. Figure S1). The morphology of these flow fields compare well with feeding flows around isolated *Vorticella* (a contractile, stalked ciliate which bears strong resemblance to *Stentor*) in the presence of a no-slip boundary [23, 24].

## 3. Ciliary coordination and emergence of metachronal waves

### 3.1. Spatiotemporal correlations

Next, we use high-speed imaging (frame rate of 1 KHz) to investigate how the arrangement and synchronicity of the membranellar cilia contributes to the restructuring of extracellular flows during oral regeneration. Image sequences were processed to localise fast-moving ciliary structures (Figure 3a and inset). Greyscale intensities measured from each region of interest were cross-correlated to determine how the spatiotemporal coordination exhibited by the beating membranelles changes over time.

**Figure 3:**
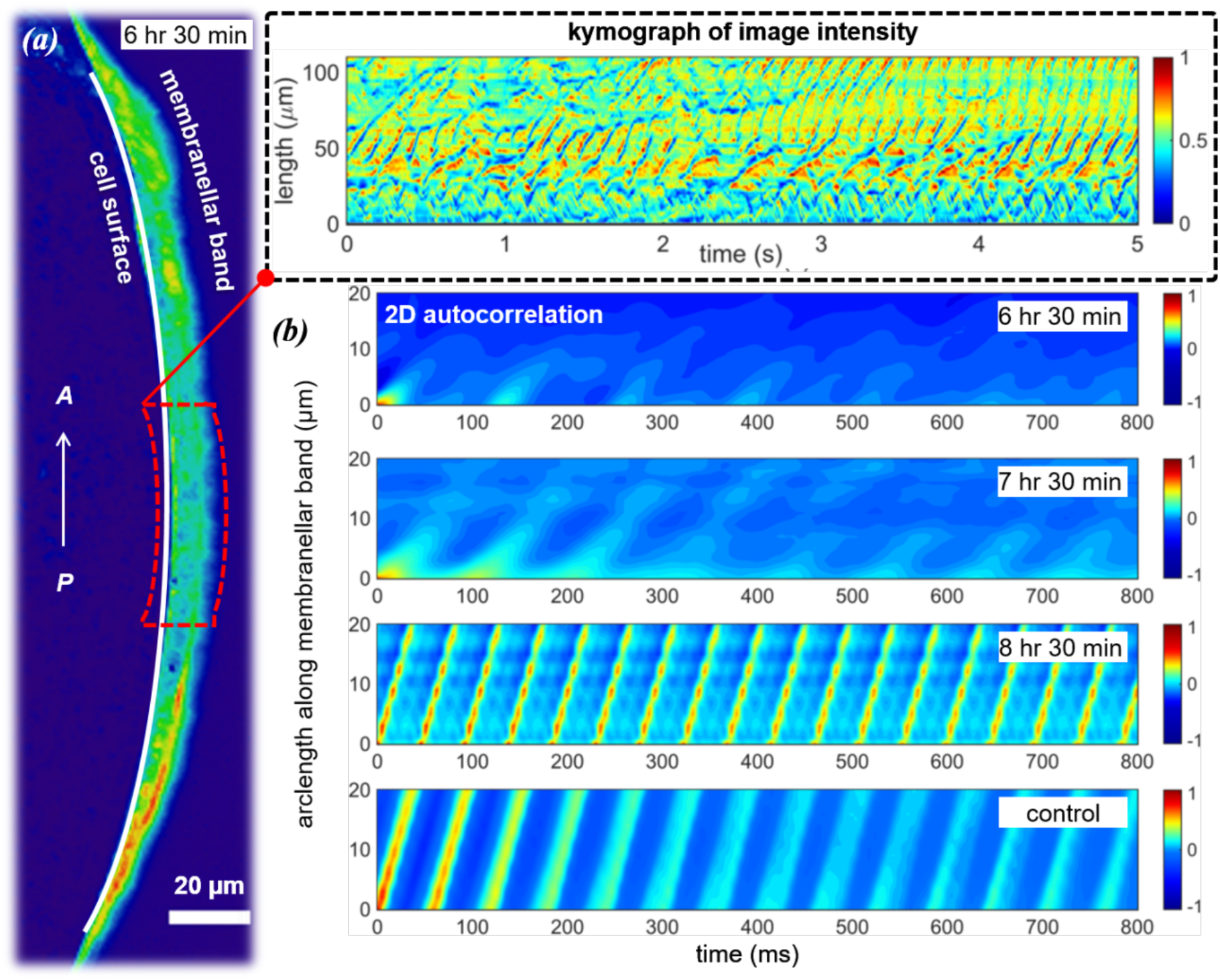
a) To measure ciliary coordination, a heatmap was used to first localise the beating cilia, e.g. to a narrow band on the surface of the organism (arrow in direction of increasing arclength from posterior to anterior). Dashed line encloses a region of interest, parallel to the ciliary band, which was used to compute intensity kymographs. Inset: kymograph can reveal local structure and coherence. b) The 2D intensity autocorrelation was used to measure changes in ciliary coordination over the course of regeneration. Sustained MCWs (parallel lines of high correlation propagating posterior-anteriorly) only emerge at very late times, here, slope = MCW speed.

At very early times, the short membranelles beat without proper coordination. Over the next 1-2 hours, the growing ciliary band remains linear and incomplete waves of activity are propagated from the newly formed oral primordium toward the anterior (Figure 3b, 6hr 30min, supp. Video 2). However, there is no long-range coordination at this stage contrary to reports in historic works [20]. Instead, characteristic wave-like structures appear and disappear intermittently along the membranellar band, but rarely extending beyond one or two wavelengths. As the reshaping structure begins to migrate toward the anterior, the MCWs become slightly more coordinated and sustained over much longer distances (Figure 3b, 7hr 30min). The individual membranelles beat in a direction perpendicular to the developing peristome to propagate a diaplectic MCW. Hereafter, the ciliary beat amplitude no longer changes, but the beat frequency and speed of wave propagation remain subject to intracellular control [25]. Very late in regeneration, a single, highly-coordinated, MC is propagated circularly from the gullet around the entire peristome - and it is only then that global coordination of the membranelles is attained (supp. Video 3). Existence of a single unidirectional wave is evident from plots of 2D intensity autocorrelations as a function of time and arclength along the developing membranellar band. In the examples shown, both control *Stentor* and very late-stage regenerating cells show parallel lines of equal slope (Figure 3b), corresponding to 0.58 mm/s and respectively 0.57 mm/s for the speed of MCW propagation, and 16.1 Hz versus 20.8 Hz for the mean beat frequencies of individual membranelles. The timing of global coordination is also coincident with the emergence of the strong feeding vortex.

### 3.2. Infraciliature and hydrodynamic interactions

To explain the lack of ciliary coordination at early times and emergence of global order at late times, we correlated our light microscopy data with transmission electron microscopy (TEM) sections of fixed *Stentor* (supp. Materials). Paulin & Bussey [20] showed previously that in the early stages of oral regeneration new membranellar cilia sprout from basal bodies that are arranged randomly in the anarchic field, consistent with observed lack of ciliary coordination. At 4-5 hrs these undergo restructuring to assemble regularly stacked rows of membranelles, with fibrillar structures connecting neighbouring membranelles also appearing at this stage (Figure 32, [6]). A rotation in direction of fluid pumping from longitudinal to transverse (Figure 2d, e) is fully consistent with the transition from a longitudinally directed beat pattern to a transversely directed beat [20]. The intermittent coordination between nearby membranelles observed at this stage (Figure 3b) can be attributed to hydrodynamic interactions: strong correlation at small spatial and temporal lags arises from strong coupling between nearby membranelles over distances of ∼10 µm, in both intermediate (cilia have already reached full-length, Figure S3) and late-stage regenerating cells. This lengthscale is comparable to the bifurcation length for hydrodynamic coupling [26]. Further evidence of hydrodynamic effects can be deduced from Tartar’s finding that ciliary coordination in grafted *Stentor* can be immediate – long before any new cellular connections could have formed between the new and old membranelles [2].

Though necessary, such passive fluid-structure interactions alone are insufficient to achieve global ciliary coordination. Instead, *Stentor* MCWs are regulated by excitatory signalling via a “silver line” system of intermembranellar connections with the primordium-facing membranelles acting as pacemaker [25]. As in some species of algal multiflagellates [27, 28], contractile elements can reverse the beating direction of the membranelles dynamically [29]. Addition of digitoxin and other chemicals can directly modulate intermembranellar connectivity leading to a change in MCW wave velocity but affect the beat frequencies to a lesser extent [30]. Existence of a single peristomal frequency must require physical continuity across entire membranellar band – indeed in non-regenerating *Stentor*, MCWs can be stopped or altered by severing or manipulating the basal fibres between membranelles [2, 31].

We conclude therefore that geometry is the final ingredient required to achieve global metachrony. In particular, by the 7 hr stage, all internal, fibrillar structures are already present (Figure 4a) as in control *Stentor*, yet coordination could not be sustained along the entire linear band of membranelles (Figure 4b). Instead, a MCW with a unique, constant wave speed only emerges after the conical reshaping of the frontal field where the entire band becomes nearly circular. These findings highlight the importance of topology [32], curvature [33], and periodic boundary conditions [34] for the stabilisation of MCWs on ciliated surfaces. Particularly, this is consistent with the notion that global ciliary coordination requires a precise orientation and localisation of stacked membranelles. We suggest that for this reason oral regeneration in *Stentor* always proceeds in the same manner in order to attain a highly-specific, ultimate ciliary configuration which can effectively exploit the interciliary hydrodynamic interactions [35, 36, 37]. Further investigation of the specialised geometric constraints exhibited by *Stentor* membranellar cilia, and their relation to the structure and dynamics of the MCWs, will appear in a follow-up study.

**Figure 4:**
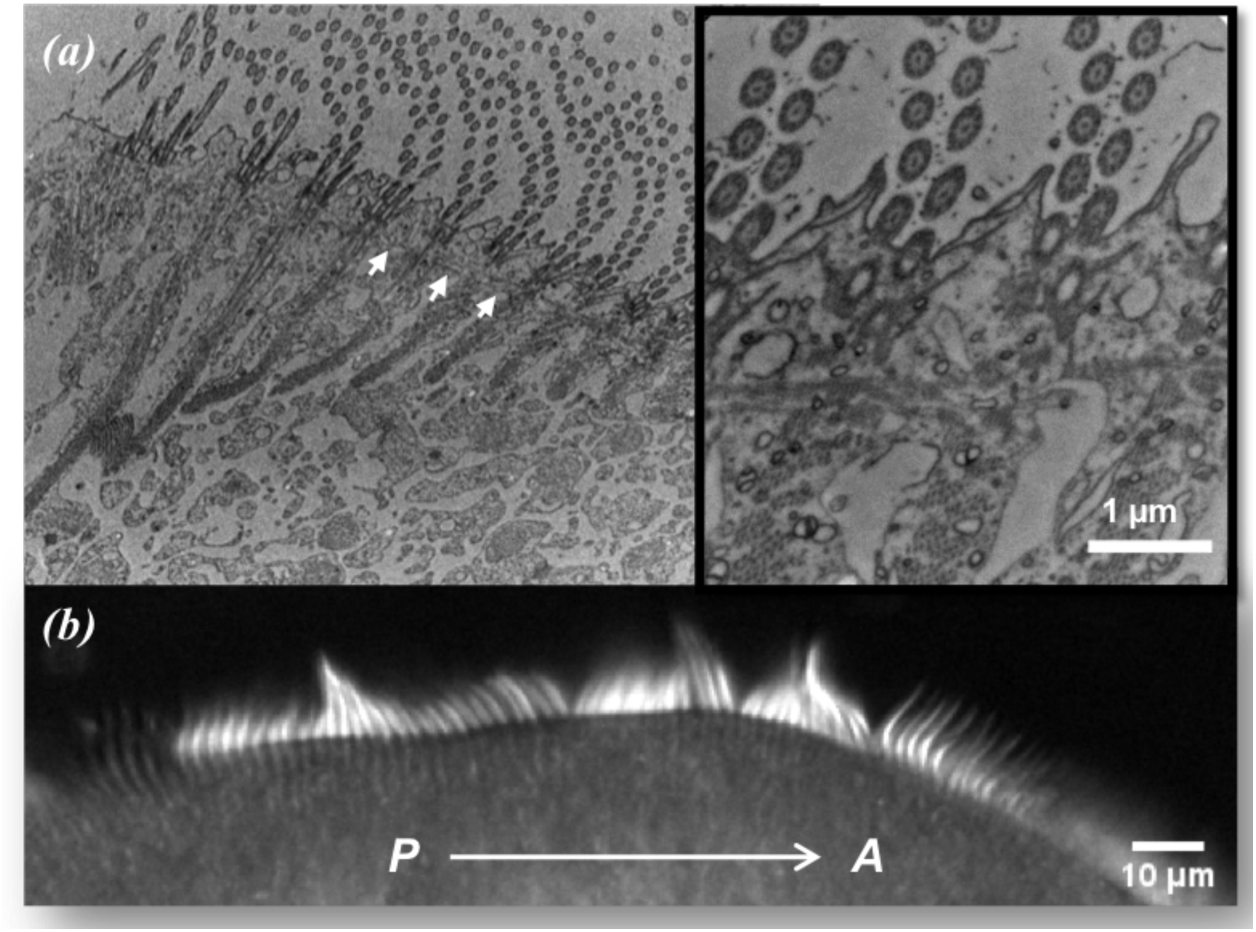
Correlating TEM sections with live-cell light microscopy imaging at the same regeneration stage -- at 6 hrs 45mins post sucrose-shock: a) the ultrastructure is indistinguishable from control *Stentor*, fibrillar structures extend from the membranelles into the cytoplasm, in addition to transverse connections between neighbouring membranelles (arrows); b) cilia at the corresponding stage in live cells exhibit local but not global coordination – transient waves are propagated from the new oral primordium situated at the posterior (P), to the anterior end of the organism (A).

## 4. Discussions and Outlook

The formation of a new oral apparatus in *Stentor* is a remarkable feat of organellar regeneration, differentiation, and structural reorganisation involving the proliferation of 15,000 individual basal bodies -- their reconfiguration and relocalisation into a functional entity, along with a precise arrangement and migration of associated subpellicular microtubules, basal-body associated fibres and other contractile filaments. Here, this process has been studied systematically in live cells for the first time with correlative flow-tracking and high-speed imaging. Live-cell imaging is indispensable when inferring dynamics and behaviour, as erroneous conclusions could be drawn from interpretation of static EM images alone. Here, we developed a robust methodology to track how spatiotemporal coherence in the membranellar structure evolves in substrate-adhered organisms over the full course of regeneration (up to 10 hours of continuous imaging), starting from incoherent random ciliary beating, to enhanced coordination and reorientation of fluid pumping, and ultimately to the emergence of a single directional MCW spanning the peristome which generates and sustains large-scale vortical flows. The synchronization dynamics of MCWs in *Stentor* add a novel dimension to a growing literature on biophysical studies of interacting arrays of cilia in existing model systems [38, 39, 28]. Specifically, the present study highlights the necessity of correct basal body and membranellar alignment, fibrillar connections, as well as hydrodynamic interactions to achieve overall metachronal coordination.

*Stentor* is an emerging model organism that has great potential for biophysical studies of ciliary regeneration and coordination, offering many novel avenues for consideration by the interested reader. Its unique and multifarious regenerative capabilities can be used in conjunction with targeted physical manipulations such as dissection, grafting or induced resorption to study ciliary and flow reorganisation. However, the novelty of this system also presents a number of challenges. Firstly, we were unable to obtain cell-cycle synchronized cultures; this coupled with the highly-contractile cell body meant that experimental organisms were highly variable in size, shape, and behaviour. Secondly even though the dynamics concerned are intrinsically three-dimensional, for constraints of phototoxicity and imaging speed we use 2D-imaging only. As part of this study, we had also intended to investigate an alternate form of oral regeneration termed *in situ* regeneration, in which membranellar cilia reputedly could be induced to sever at the transition zone and regrown in place (as in the biflagellate alga *Chlamydomonas* [40]), without any morphological restructuring of the cytoskeleton [41, 42]. *In situ* regeneration would have enabled the effective decoupling of intracellular or ultrastructural changes from passive hydrodynamic interactions during cilia and membranellar regrowth, but after multiple attempts the present authors were unable to reproduce this configuration - the membranellar cilia were always shed together with the entire membranellar band. Finally, we suggest that microsurgical manipulation can provide further insights into the ciliary coordination mechanism.

Despite these sources of variability, our results demonstrate a remarkable consistency in the timing and dynamics of *Stentor* oral regeneration. Electing to use the sucrose-shock method we were able to induce reversible and simultaneous shedding of membranellar bands in large sample sizes. Physiological regeneration, for instance due to injury of feeding organelles or excess mechanical stresses, also elicit a similar programmed response [43, 4]. Thus, membranellar regeneration appears to be a universal mechanism by which unicellular protists repair and refresh their oral structures [44]. The concomitant change in topology from linear to circular appears to be critical for attainment of global coordination of membranelles and the inward-directed feeding flows. In this regard, protists such as *Stentor* that can achieve great complexity of cortical organisation and internal control over locomotor appendages, could hold the key to understanding the evolution of neural circuitry [45, 46] designed for deriving coherence of ciliary arrays.

## Endnotes

### Authors’ Contributions

KYW & SKH conceived the study, performed regeneration experiments, data analysis, and wrote the manuscript; AMF performed IF experiments; RMM & TM cultured organisms, assisted with microscopy, contributed EM data; EB & JYS obtained flow data for unattached organisms; WFM secured funding, provided supervision, and initiated the collaboration at the MBL in Woods Hole. All authors contributed to project design and article revision, and have given approval for the final version.

### Funding

We gratefully acknowledge financial support from the Marine Biological Laboratory at Woods Hole, MA, NIH grant R35 GM097017 (WFM) and the University of Exeter, UK (KYW).

**The following multimedia files accompany this study:**

1. DIC imaging of flow fields around a regenerating *Stentor* (5.5 hrs post sucrose shock)
2. High-speed video of linear band (6.5 hrs post sucrose shock)
3. High-speed video of fully-regenerated membranellar band (8.5 hrs post sucrose shock)

## Acknowledgments

We thank Greyson Lewis and Akanksha Thawani for discussions. We thank Kasia Hammer (Central Microscopy Facility, MBL) for preparing samples for electronmicroscopy, and the Nikon Imaging Center at UCSF for access to imaging facilities. We remain indebted to the staff and students of the MBL Physiology Course for creating a stimulating environment for scientific research.

